# Conserved +1 translational frameshifting in the *S. cerevisiae* gene encoding YPL034W

**DOI:** 10.1101/2020.04.29.069534

**Authors:** Ivaylo P. Ivanov, Swati Gaikwad, Alan G. Hinnebusch, Thomas E. Dever

## Abstract

Living cells have developed exquisite mechanisms to ensure accurate translation of mRNA. Many of them are dedicated to preventing the change in reading frame during translation elongation. A minority of chromosomally encoded genes have evolved sequences that subvert standard decoding to program +1 translational frameshifting, either constitutively or in response to external stimuli. In the yeast *Saccharomyces cerevisiae*, three chromosomal genes are known to employ programmed +1 translational frameshifting for expression of full-length functional products. Here we identify a fourth yeast gene, *YFS1*, encompassing the existing predicted open reading frame *YPL034W*, with conserved programmed +1 frameshifting. Like the previously known examples, it appears to exploit peculiarities in tRNA abundance in *S. cerevisiae*.

## Introduction

As determined more than half a century ago, translation begins with ribosomes initiating at a specific start codon, usually AUG, which commits them to translating one of the three possible reading frames of an mRNA (Adams and Capecchi 1966). Elongation then proceeds by decoding adjacent triplet nucleotides until one of three stop codons is encountered (Crick et al. 1961). Loss of frame maintenance has severe consequences, resulting in loss of protein expression leading to haploinsufficiency (Morrill and Amon 2019) or translation of aberrant peptides that also terminate prematurely or with C-terminal extensions leading to dominant negative genetic consequences or activation of cellular stress responses (Herskowitz 1987; Ince et al. 1993; Sham et al. 2001; Wang and Kaufman 2012). As a result, cells invest considerable energy and resources to frame maintenance once ribosomes have committed to the start of translation. Nevertheless, a small group of genes have evolved sequences that act in *cis*-to subvert standard decoding, often for regulatory purposes – a process known as recoding (Atkins and Gesteland 2010). Recoding encompasses phenomena like stop codon readthrough, ribosome hopping, and ribosomal frameshifting. In the latter, a fraction of translocating ribosomes slip forward (5’) or backward (3’) resulting in “+” or “-” frameshifting (Farabaugh 1996; Atkins et al. 2016). Most known cases of programmed frameshifting involve either +1 or −1 frameshifts that, while superficially similar, are mechanistically distinct. Well studied examples of −1 frameshifting result from simultaneous slippage of tRNAs relative to mRNA when both the A and P sites of the ribosome are occupied by tRNAs decoding a heptanucleotide sequence with the signature motif X-XXY-YYZ (Jacks et al. 1988), wherein the tRNA decoding the triplet XXY can also decode XXX and that decoding YYZ can also decode YYY. By contrast, +1 frameshifting usually occurs when the ribosome P site is occupied by peptidyl-tRNA, but the A site is empty owing to slow decoding of the A site triplet (Weiss et al. 1987; Hansen et al. 2003). In addition to a defined frameshift site sequence, most cases of programmed frameshifting include additional sequences, present 5’ or 3’ of the shift site, that can dramatically alter the efficiency of frameshifting (Atkins et al. 2016).

Work on translation of the gag-pol fusion proteins encoded by the Ty1 and Ty3 retrotransposons of *S. cerevisiae* provided important insights into the mechanism of +1 frameshifting. Ty1 frameshifting occurs on the heptanucleotide sequence CUU-AGG-C when the CUU Leu codon is in the ribosome P site and AGG Arg codon is in the A site (Belcourt and Farabaugh 1990). This sequence alone, without any additional *cis*-elements, can support 40% or more frameshifting depending on the reporter. A larger cassette, including additional sequences naturally surrounding the shift site, gives somewhat lower +1 frameshifting of 12-20%, suggesting the potential presence of frameshift suppressing sequences (Clare et al. 1988; Harger and Dinman 2003). The CUU-AGG-C sequence also directs +1 frameshifting in *ABP140* (Asakura et al. 1998), one of three *S. cerevisiae* chromosomal genes known to employ programmed +1 frameshifting. Another yeast gene requiring +1 frameshifting for expression, *EST3*, employs a derivative of the Ty1shift site, CUU-AGU-U (Morris and Lundblad 1997). The third *S. cerevisiae* chromosomal gene, *OAZ1*, has the shift site GCG-UGA-C (Palanimurugan et al. 2004), which shares a P-site codon with the Ty3 shift site, GCG-AGU-U (Farabaugh et al. 1993). The latter, in turn, shares the A-site triplet with that of the *EST3* shift site. If the shift prone sequence CUU-AGG-C is present in-frame within a main Open Reading Frame (mORF), which otherwise does not require +1 frameshifting, it will result in a high frequency of spurious switches in the reading frame, producing truncated polypeptides, which could be detrimental to cell physiology. Computational analysis supports this prediction by showing significant underrepresentation of the heptanucleotide CUU-AGG-C in-frame in yeast mORFs, even when codon usage and nucleotide composition are taken into account (Shah et al. 2002).

Here we show that the previously annotated yeast ORF *YPL034W* is part of a larger conserved ORF lacking an initiation codon that is instead accessed following initiation at an overlapping ORF initiated further upstream. A fraction of ribosomes initiating at the first ORF frameshift in the +1 direction while translating the heptanucleotide sequence CUU-AGG-C. This frameshift sequence, and the frameshifting itself, is conserved in most sequenced members of the budding yeast *Saccharomycetaceae* family. We propose to name the newly identified gene Yeast Frame Shift 1 (*YFS1*).

## Results

### Evidence that expression of *YPL034W* requires +1 frameshifting

During a search for conserved upstream open reading frames (uORFs) in yeast, we visually examined the mRNA of *YPL034W* for its translation potential using global aggregate ribosome profiling data available at GWIPS-viz (Michel et al. 2014). Aggregate 80S ribosome profiling coverage is consistent with translation initiating at an AUG codon at positions 486,411-486,413 of chrXVI (SacCer_Apr2011/sacCer3 assembly) located 298 nucleotides (nt) upstream of the annotated AUG start codon of *YPL034W* (at positions 486,712-486,714) indicated at SGD (https://www.yeastgenome.org). Ribosome footprints proximal to this upstream AUG codon are denser than the footprints in the annotated mORF of *YPL034W* (Fig. 1A), which led us initially to consider the AUG as the start codon of a short uORF of 20 codons. Surprisingly, however, there is no break in 80S ribosome coverage between the stop codon of the suspected uORF and the annotated mORF beginning 239 nt downstream. This is true even after taking into account the presence of a potential additional uORF of 111 nt starting 30 nt downstream of the first suspected uORF (Fig. 1A, B). The lack of any change in ribosome coverage in the region before and after the annotated AUG start codon of *YPL034W* suggested the possibility that this region is translated without interruption starting somewhere within the suspected first short uORF.

**Figure 1.**
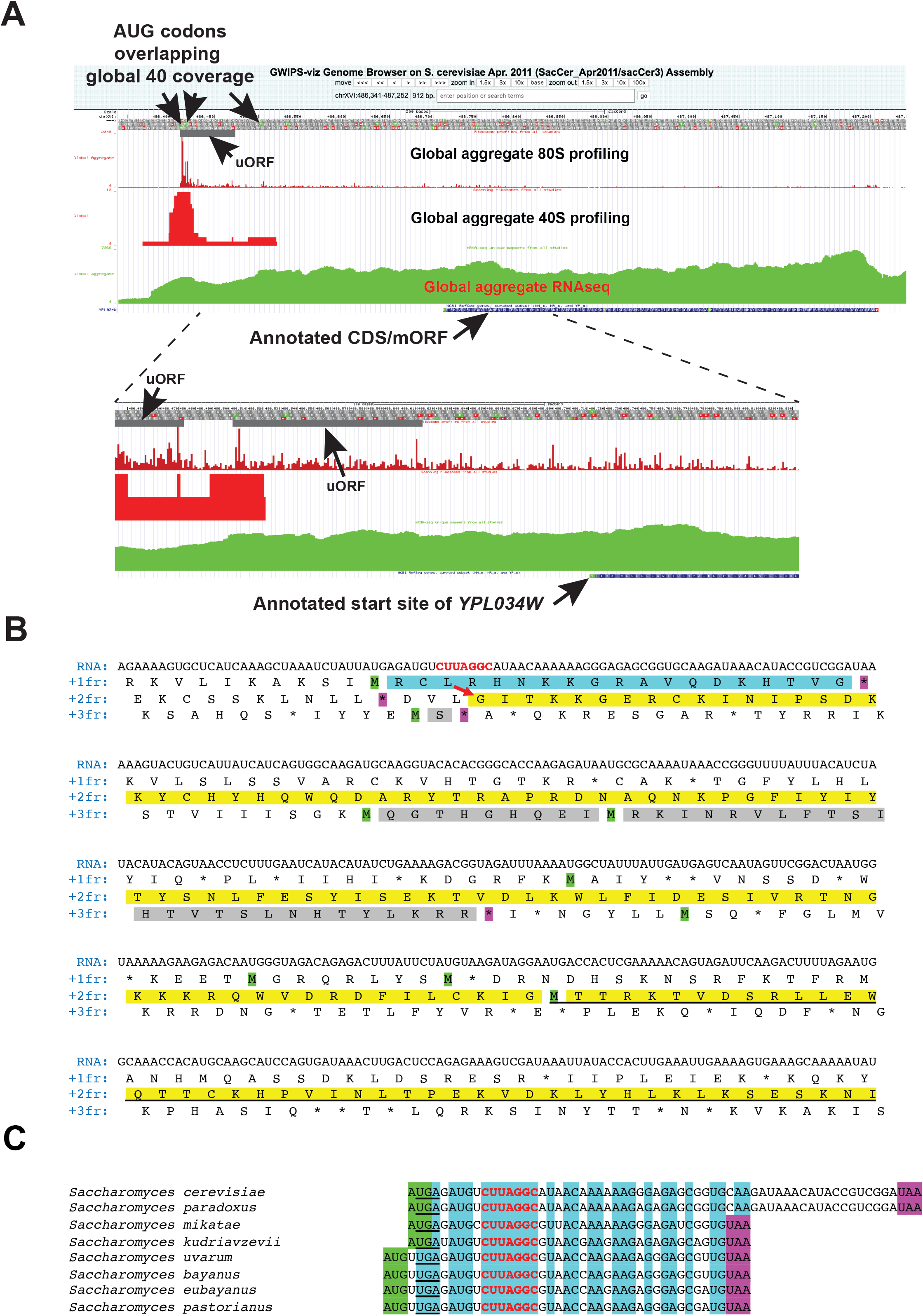
Aggregate ribosome profiling and the structure of the mRNA encoding YPL034W provides evidence for ribosome frameshifting. (A) Global aggregate data for 80S, 40S and RNAseq coverage – screenshot of the vicinity of *YPL034W* as displayed in GWIPS-vis (https://gwips.ucc.ie/). The amino acid sequence in all three reading frames of the positive strand is shown as a band near the top (the exact sequence can be seen in (B)). AUG start codons are in green, and stop codons are in red. The position of the first uORF is indicated; and the global 80S coverage and aggregate 40S coverage are displayed below. The positions of AUG codons corresponding to global aggregate 40S coverage are indicated by arrows pointing down. Underneath the 40S coverage is aggregate RNA coverage. The position and sequence of the annotated *YPL034W* ORF is shown below the RNA coverage. Lower panels: Blow up of the region spanning the first uORF to the annotated mORF, with the position of a second putative uORF also indicated. (B) Nucleotide and amino acid sequence of approximately the first 2/3 of *S. cerevisiae YFS1* mRNA. The sequence begins with the transcription start site as determined by CAGE data (Lu and Lin 2019). The mRNA nucleotide sequence is on top, and the simulated translated amino acid sequence in all three frames is shown below it, with the different frames indicated on the left. Methionine residues, encoded by putative initiation AUG codons, 5’ of the annotated AUG start codon of *YPL034W* are highlighted in green. Relevant in-frame stop codons are in magenta. The sequence of the first uORF of *YPL034W* (ORF1 of *YFS1*) is highlighted in cyan. The proposed encoded peptide of ORF2 of *YFS1* is highlighted in yellow; the part constituting the previously annotated *YPL034W* is underlined. Two uORFs/iORFs discussed in the main text are highlighted in gray. The putative frameshift site of *YFS1* in ORF1 is shown in red letters. The position and direction of the putative +1 translational frameshift is indicated by a red arrow. (C) Nucleotide sequence alignment of ORF1 (annotated uORF) of *YFS1*. The sequences of ORF1 from 8 species belonging to the *Saccharomyces* genus are aligned on the right. Species names are on the left. Start codons are highlighted in green. Stop codons are highlighted in magenta. Nucleotides conserved in all 8 species are highlighted in cyan. Nucleotides constituting the putative frameshift site are in red. The in-frame UGA stop codon bracketing the 5’ end of ORF2 in the +1 frame is underlined.

Performing 40S ribosome profiling Archer et al. previously demonstrated accumulation of 40S footprints at annotated initiation codons and depletion of 40S footprints immediately downstream (Archer et al. 2016). The GWIPS-viz profile of this 40S profiling data are consistent with initiation at the AUG start codon of the suspected first uORF and possibly at the AUG start codon of the second potential uORF beginning 30 nt further downstream (Fig. 1A, B). The 40S data do not, however, provide support for initiation at the annotated AUG start codon of the mORF. Mapping of transcription start sites (TSSs) by Cap Analysis of Gene Expression (CAGE) revealed that the major cluster of TSSs found at *YPL034W* maps ∼37 nt upstream of the first AUG codon in the mRNA (the start codon of the potential first uORF (http://yeastss.org/). Together, these data support the hypothesis that the AUG start codon of the potential uORF1 is actually the primary start codon of *YPL034W*.

The sequence of the first 465 nt of the *S. cerevisiae YPL034W* encoding mRNA and the features identified in it are shown in Fig. 1B. The coding region corresponding to the annotated (coding sequence) CDS (here also referred to as the mORF) in *S. cerevisiae* can be extended upstream without intervening in-frame stop codons to just 3’ of the start codon of the first uORF, which is in a different frame (Fig. 1B). Sequences from 7 additional homologs of *YPL034W* from yeast belonging to the *Saccharomyces* genus were obtained for analysis. In all 7 sequences, the bracketing (upstream) in-frame stop codon for the mORF is also just after the start codon of the first uORF, which too is conserved in all 8 *Saccharomyces* species (underlined in Fig. 1C). No available in-frame AUG codons exist to initiate translation of this extension of the mORF, as is also true for all analyzed *Saccharomyces* orthologs. For all 8 sequences, the reading frame corresponding to the annotated mORF was conceptually translated from the first in-frame stop upstream of the annotated start codon to the next in-frame stop codon (the annotated stop codon of the mORF). The sequences were then aligned using ClustalX and displayed as a logogram (Fig. 2A). The alignment shows a number of amino acid residues 5’ of the annotated AUG start codon that are conserved in all 8 species, consistent with the idea that this region is translated and that its product has biological function. We considered the possibility that the conserved extension is initiated by a near-cognate start codon, as has been reported for various genes in both in *S. cerevisiae* and in other species (Peabody 1989; Chang and Wang 2004; Tang et al. 2004; Ivanov et al. 2011; Wei et al. 2013). However, no conserved near-cognate start codon is apparent 3’ of the upstream in-frame stop codon and 5’ of the first absolutely conserved amino acid residue, T6, seemingly precluding the possibility of near-cognate initiation of the extension.

**Figure 2.**
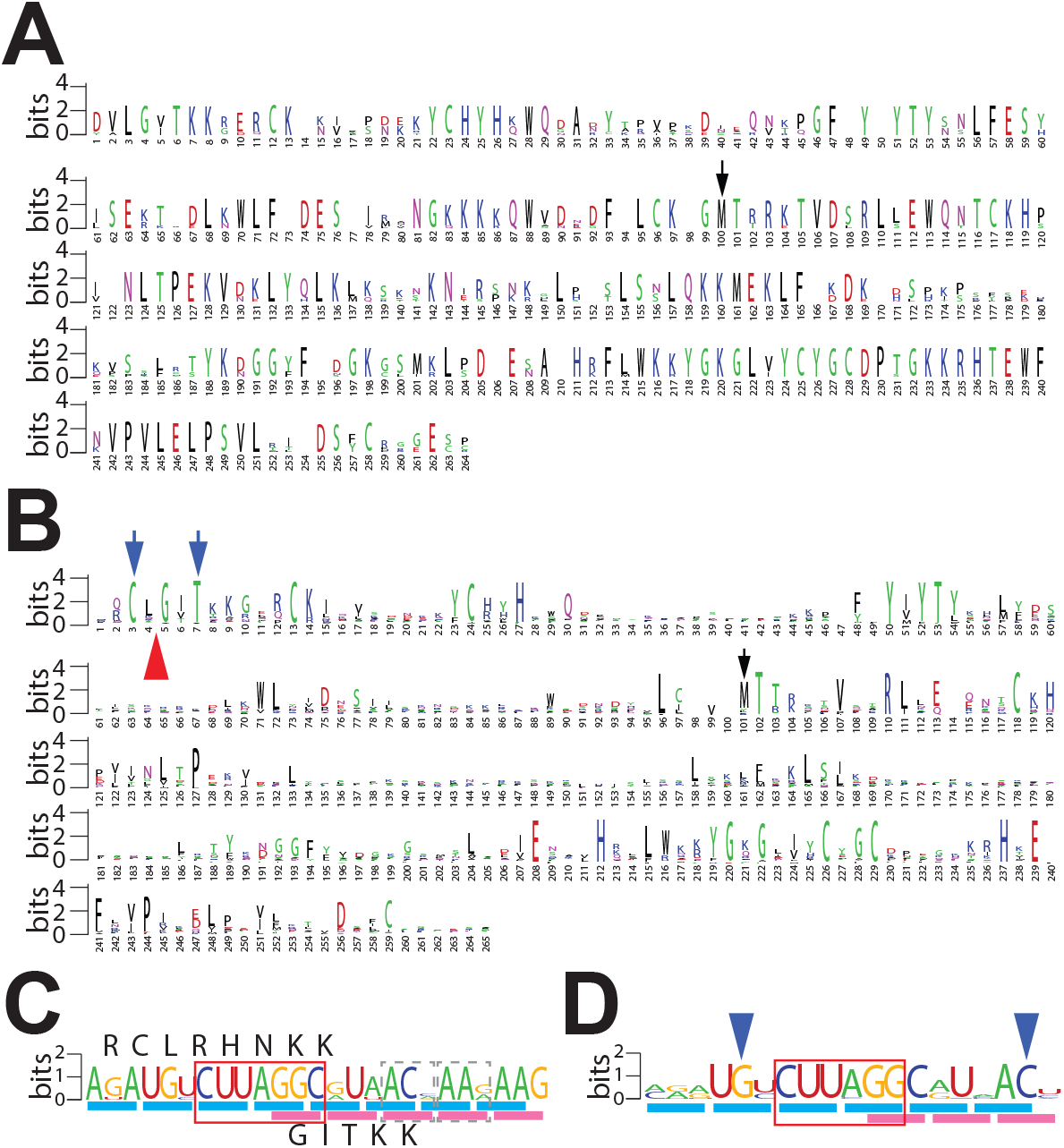
Conservation of the amino acid sequence of Yfs1 upstream of the annotated YPL034W and of the nucleotide sequence of the frameshift site. (A-B) Weblogo representation of the amino acid sequence conservation based on (A) alignment of ORF sequences (upstream in-frame stop codon to annotated stop codon of YPL034W) from 8 species belonging to the genus *Saccharomyces* or (B) alignment of simulated translations of sequences from 81 species belonging to the fungal order *Saccharomycetales* – 52 with and 29 without a frameshifting site. Insertions relative to the *S. cerevisiae* sequence were removed, and therefore, numbering corresponds to the *S. cerevisiae* sequence. In both (A) and (B) the position of the currently annotated AUG (YPL034W) start codon is indicated by a black arrow. In (B) the position of the frameshift site is indicated by a red arrow; the position of the two conserved codons flanking the frameshift site (shown in D) are indicated by blue arrows. (C-D) Weblogo representation of the nucleotide sequence conservation flanking the frameshift site of *YFS1* based on (C) sequences from 8 species belonging to the genus *Saccharomyces* or based on (D) the same sequences from 81 species belonging to the fungal order *Saccharomycetales* indicated in (B). Triplets in the “0” frame are underlined with blue bars, while those in the “+1” frame are underlined in magenta. The CUU-AGG-C Ty1-like shift site is boxed in red. In (C) two triplets showing synonymous substitution pattern in the +1 frame following the frameshift site are boxed with dashed lines. The amino acid sequence in the 0 and +1 frame in *S. cerevisiae* are shown above and below the conservation logo, respectively. In (D) triplets (UGY Cys and ACY Thr) showing synonymous substitution patterns before and after the shift site, in the “0” and “+1” frame, respectively, are indicated by blue arrows.

We next focused our attention on the first potential uORF. In all 8 species, this uORF overlaps the putative N-terminal extension of the *YPL034W* mORF in the +1 frame. However, even in these relatively closely related species, neither the position of its start nor its stop codon is invariant (Fig. 1C). Aligning the nucleotide sequences of the predicted first uORF for the eight *Saccharomyces* homologs of *YPL034W* (Fig. 1C) immediately suggested a mechanism for the translation of the putative N-terminal extension. Seven absolutely conserved nucleotides, CUU-AGG-C, were found within the potential uORF. Of note, the CUU triplet is in-frame with the uORF AUG start codon; moreover, in the +1 reading frame of the motif (the frame of the mORF) there are no stop codons between the motif and the annotated mORF start codon (Fig. 1B). Strikingly, this conserved sequence is identical to the known +1 frameshift site required for translation of gag-pol in the retrotransposon Ty1, and also in the yeast chromosomal gene *ABP140*. As noted above, a variant of this sequence, CUU-AGU-U, is responsible for the +1 frameshifting in the yeast chromosomal gene *EST3*. Furthermore, almost immediately following the motif (putative shift site) in *YPL034W*, the pattern of nucleotide conservation is consistent with synonymous substitutions in the +1 frame, i.e. triplets ACN (Thr) and AAR (Lys) (Fig. 2C - boxed in grey). Together, these findings strongly suggest that translation of the mORF of *YPL034W* proceeds by initiation at the AUG start codon of the first potential uORF and then shifts into the continuous reading frame of the mORF owing to slow decoding of the AGG triplet with subsequent decoding of the GGC triplet instead following a +1 frameshift. Henceforth, to distinguish between the newly identified gene product and the previously annotated (partial) *YPL034W* ORF, we refer to the full-length protein product as Yeast Frame Shift 1, Yfs1, and the corresponding gene *YFS1*. The first predicted uORF will be referred to as ORF1 and the N-terminally extended mORF of *YPL034W* will be designated ORF2.

### Natural history of the frameshift site

To determine how widespread frameshifting in *YFS1* might be among yeast species, an additional 73 sequences of *YFS1* homologs from the yeast order *Saccharomycetales* were retrieved, for a total of 81 homologs (see Materials and Methods). The sequences were manually examined for their potential, or requirement, for +1 frameshifting, conceptually translated, and the predicted peptide sequences subjected to additional analysis. Of the 81 homologs, 52 require +1 frameshifting for expression of the protein, while 29 do not. ClustalX was used to align the predicted amino acid sequences of all 81 proteins and to generate a phylogenetic tree based on this alignment. The alignment (Fig. 2B) shows that even though the peptide sequence is diverging at a relatively high rate, a number of amino acid residues in the predicted N-terminal extension of ORF2 appear to be as highly conserved as do certain residues located downstream of the methionine corresponding to the annotated start codon of *YPL034W* (amino acid 101 in the alignment, marked with a black arrow in Fig. 2A, B), presumably because they are crucial for the function of the protein.

Alignment of the nucleotide sequences surrounding the putative frameshift site from the 52 homologs that require +1 frameshifting shows near-perfect conservation of the CUU-AGG-C heptanucleotide sequence (Fig. 2D). In three related species, the heptanucleotide sequence is CUU-CGG-C instead. In addition to the heptanucleotide shift site, several additional nucleotides are highly conserved – a “UG” dinucleotide before the shift site and a “UNAC” motif located downstream. Further analysis of the amino acid sequence conservation revealed that the Cys specified by codon 3 of *S. cerevisiae YFS1* (encoded by UGY) is absolutely conserved in all 81 sequences examined, including 29 that do not require frameshifting, and is therefore likely essential for the function of the protein. This indicates that the UG dinucleotide before the frameshift site is likely conserved because of the importance of the amino acid it encodes rather than its role in frameshifting. The same is probably true for the Thr (ACY) codon at position 7 of the protein that includes the AC dinucleotide of the UNAC motif. The only nucleotide outside the putative heptanucleotide sequence that appears conserved because of a likely role in frameshifting is the “U” of the UNAC motif two nucleotides downstream of the putative heptanucleotide shift site, as it occurs as the 2^nd^ base in a variety of triplets encoding multiple different amino acids.

The phylogenetic tree of the 81 *YFS1* homologs matches the known phylogenetic relationship (https://www.ncbi.nlm.nih.gov/Taxonomy/) of the corresponding yeast species (Fig. 3). All species that branch prior to the split between the family *Saccharomycetaceae* and the other members of the order *Saccharomycetales* lack a requirement for frameshifting. Members of the genus *Kluyveromyces*, which branched off early during the diversification of *Saccharomycetaceae*, also do not require +1 frameshifting for expression of *YFS1*. This places the emergence of +1 frameshifting in *YFS1* between 100 and 150 million years ago (Langkjaer et al. 2003; Farabaugh et al. 2006; Correia et al. 2019). Five species in the *Kazachstania* genus appear to have sustained secondary loss of +1 frameshifting. In addition, three other species in the same genus have evolved a variant of the shift site, CUU-CGG-C in which the first nucleotide of the triplet occupying the A site in the zero frame is changed from A to C. Changing the Ty1 shift site sequence from the wild type CUU-AGG-C to CUU-CGG-C in *S. cerevisiae* severely reduces the level of +1 frameshifting, but does not eliminate it completely (Belcourt and Farabaugh 1990).

**Figure 3.**
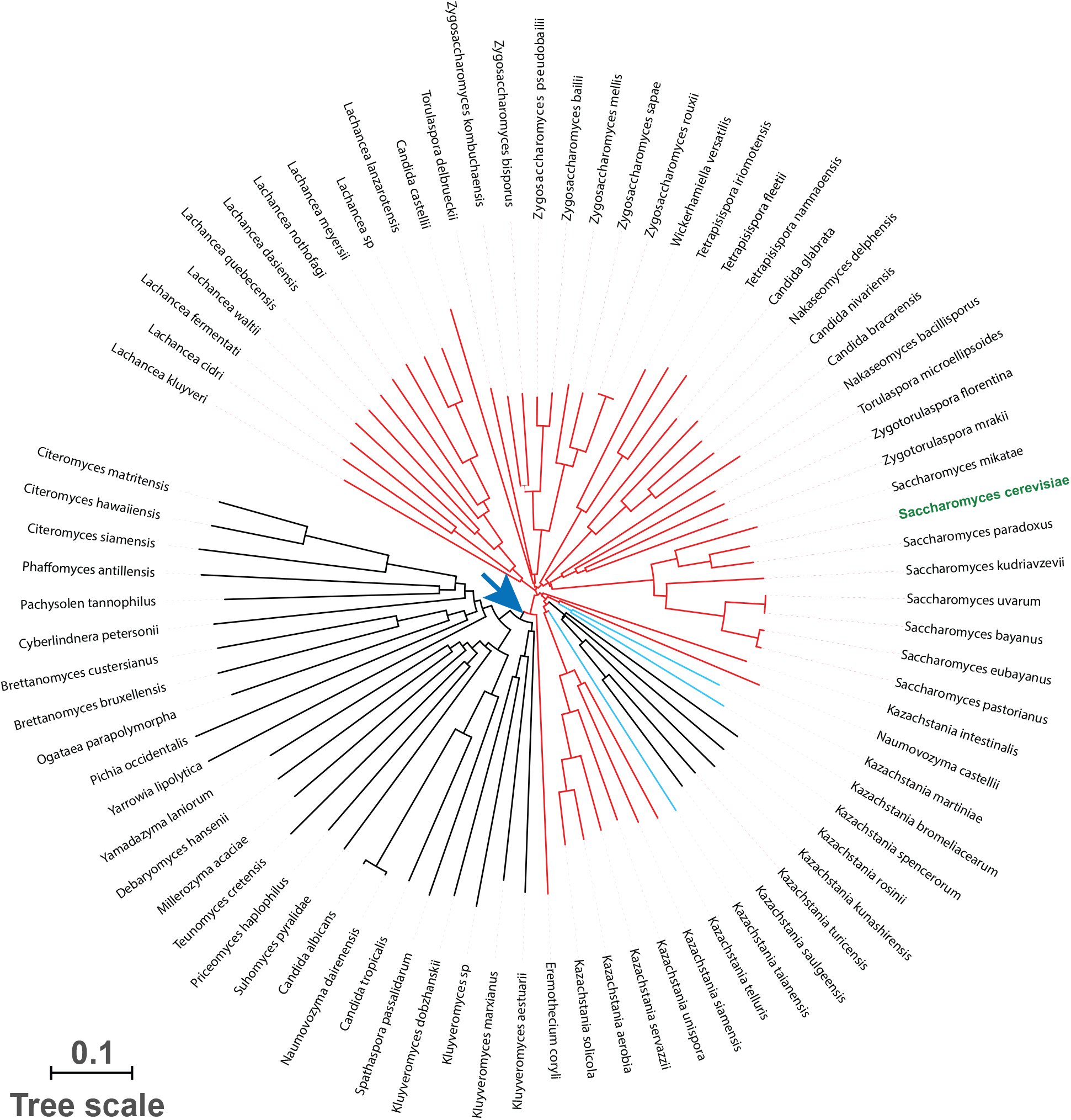
The +1 frameshifting of *YFS1* is ancient. Unrooted phylogenetic tree of 81 homologs of Yfs1/YPL034W based on their peptide sequence (after simulated +1 frameshifting) and using ClustalX. Species names are indicated on the periphery. *S. cerevisiae* is in green. The branches of homologs that require +1 frameshifting and have the inferred shift site CUU-AGG-C are in red. The branches of homologs with the variant shift site CUU-CGG-C are in blue. The branches of homologs not requiring frameshifting are in black. An inferred point of frame shift site emergence is indicated by a blue arrow. The scale bar in the lower left measures the number of amino acid substitutions per site (accounting for multiple substitutions at the same site).

### Ribosome profiling data provide direct support for +1 frameshifting in *YFS1*

Certain ribosome profiling protocols produce exceptional reading frame phase information, especially if analysis is restricted to protected fragments with a certain fixed length and direction of offset (Kiniry et al. 2019). For example, in some yeast studies more than 90% of fragments that are 28 nt long show the same 15 nt distance from the 5’ end of the footprint to the first nucleotide of the A-site codon within the ribosome. We analyzed data from several previously published studies that match these criteria and also have high ribosomal coverage (see Materials and Methods). These data were used to interrogate the evidence for +1 frameshifting in ORF1 of *YFS1*. To validate the approach, we first examined the evidence for the well-studied example of +1 frameshifting during translation of the *gag-pol* gene of yeast Ty1, which utilizes the same frameshift site, CUU-AGG-C, as the putative site in *YFS1*. In total, more than 1.3 million ribosome footprints from these studies align to the CDS of the Ty1 *gag-pol* mRNA. Plotting the reads corresponding to each frame in different colors, zero frame in blue and +1 frame in orange, a sharp change in frame signal is seen at the point of the known frameshifting site (Fig. 4A *i*). Before the frameshift site 91% of reads align at their 5’ end to the arbitrarily defined frame 1, corresponding to frame 0 of *gag*, but after that point 92% of reads align to frame 2, corresponding to frame +1 relative to *gag* and frame zero relative to *pol*. Dividing the riboseq coverage of protected fragments (normalized to total number of codons) coming from frame zero of *pol* (orange in Fig. 4A *i-ii*) by those coming from the zero frame of *gag* (blue in Fig. 4A *i-ii*) allows us to estimate the frequency of frameshifting in *gag-pol* as 10.7%, which is close to the 12% Ty1 frameshifting measured with reporters (Harger and Dinman 2003). Higher frameshifting efficiency of 20% has also been reported using a different reporter system (Clare et al. 1988). The frameshifting is even higher, 40%, with a minimal sequence including only the heptanucleotide CUU-AGG-C (Belcourt and Farabaugh 1990). The high coverage of ribosome protected fragments for this mRNA, with average reads of over 2000 per codon for ORF1, allows for near nucleotide resolution of translating ribosome positions on this mRNA. As a result, the switch of reading frames can be visualized directly (Fig. 4A *iii*), where a fraction of ribosomes can be seen moving from the “AGG” (Arg) codon in the zero frame (blue) to “GGC” (Gly) in the +1 frame (orange), with the remainder continuing in the zero frame (blue) (Fig. 4A *iii*).

**Figure 4.**
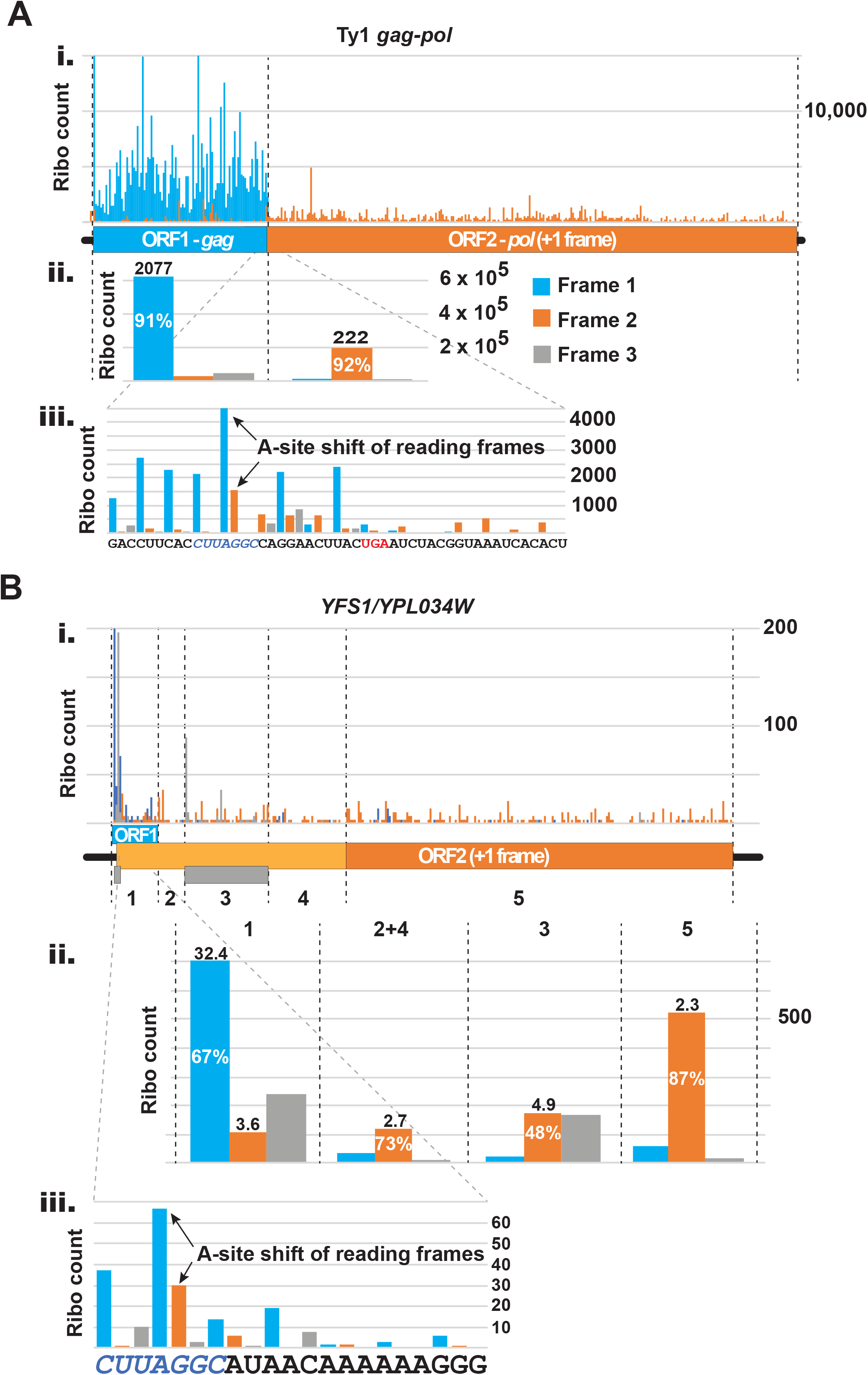
High framing quality ribosome profiling data are consistent with +1 frameshifting on the CUU-AGG-C sequence of *YFS1*. Ribosome profiling data from selected publicly available studies with high quality framing information (see main text for criteria used, methods of analysis, and sources of the sequencing data) were analyzed for evidence of frameshifting on the CUU-AGG-C sequence of Ty1 *gag-pol* and *YFS1*. Reads (28 nt only) of protected fragments were aligned at the 5’ ends with 15 nt offset to the ribosome A site. Fragments in blue align to nucleotide “1” of triplets in the “0” reading frame; fragments in orange align to nucleotide “2” of the “0” reading frame (and nucleotide “1” of triplets in the “+1” reading frame),; and fragments in grey align to nucleotide “3” of the “0” reading frame. (A) Protected fragment alignment to the mRNA of Ty1 *gag-pol*. The regions of ORF1 (*gag*) and ORF2 (*pol*) are delimited by dashed lines. Global coverage is shown on top (*i*). Middle panel (*ii*), the reads of ORF1 and ORF2 are added and tabulated by sub-codon position. The percent of total reads are written on the columns; numbers above columns are the average reads for that nucleotide position per codon. Bottom panel (*iii*), a blow-up version of the shift site is shown. Paused ribosomes with either “AGG” (before shift) or “GGC” (after +1 shift) in the A site are indicated by arrows. The corresponding nucleotide sequence of the mRNA is shown at the bottom with the shift site CUU-AGG-C in blue and in italics, and the UGA stop codon of ORF1 in red. (B) Protected fragment alignment to the mRNA of *YFS1* (*YPL034W*). Raw read counts are shown above a schematic of the mRNA features (*i*). ORF1 is in blue; the previously annotated portion of ORF2 (i.e. *YPL034W*) is in dark orange, and the 5’ extension of ORF2 is in light orange; two short internal ORFs (iORFs) in the third frame, which exhibit evidence of translation, are shown as gray rectangles. For analysis purposes, the mRNA is subdivided into different regions numbered 1-5 and the boundaries are shown as dashed lines. Region “1” = ORF1; regions 2 and 4 = sections of ORF2 that are currently not annotated as coding and do not overlap with iORF2; region 3 = iORF2; region 5 = currently annotated *YPL034W* mORF. Middle panel (*ii*), the reads aligning to the numbered regions are added and tabulated by sub-codon position. Numbers in and on top of the columns represent the same as in (A). Bottom panel (*iii*), a blow-up version of the shift site is shown. Paused ribosomes with either “AGG” (before shift) or “GGC” (after +1 shift) in the A site are indicated by arrows. The corresponding nucleotide sequence of the mRNA is shown at the bottom with the shift site CUU-AGG-C in blue and italics.

The same approach used to validate the technique on Ty1 *gag-pol* frameshifting was applied to investigate the putative frameshifting of *YFS1* (Fig. 4B). Expression levels of *YFS1* are almost three orders of magnitude lower than those for Ty1 *gag-pol*, which makes the results less unambiguous. The picture is also complicated by the presence of two apparently translated internal ORFs) encoded in different reading frames within ORF1 or ORF2 (iORFs, grey boxes in Fig. 4B *i*). Despite these complications, the data show that 67% of footprints coming from ORF1 of *YFS1* (region 1 of Fig. 4B *i-ii*) align to frame 1 (frame 0 of ORF1). Seventy-three percent of fragments aligning to the region of ORF2 that extends 5’ of the annotated AUG start, and does not overlap with the second iORF (regions 2 and 4 of Fig. 4B *i-ii*), align to frame 2 (frame 0 of ORF2), similar to the 87% of fragments that align to frame 2 (frame 0 of ORF2) of the previously annotated section of *YFS1* (region 5 of Fig. 4B *i-ii*). Evidence also exists for translation of the two iORFs, both in frame 3, one contained wholly within ORF1 (the first predicted uORF), and the second predicted uORF (now considered an iORF because it lies within ORF2 of *YFS1*) starting 30 nt downstream of the first uORF. Similar to Ty1, and despite the lower coverage, the change of frame in the A-site codon of translating ribosomes can be visualized by the aggregate 80S ribosome profiling data (Fig. 4B). By dividing the density of footprints coming from frame zero of ORF2 to the density of footprints coming from the sum of frame zero and frame +1 of ORF1 we can estimate the rate of +1 frameshifting as ∼19%. Because of the uneven and relatively limited coverage of both ORF1 and ORF2, the accuracy of this estimate is also limited. Just like with the Ty1 frameshift site, the switch of reading frames can be visualized directly, where a fraction of ribosomes can be seen moving from the “AGG” (Arg) codon in the zero frame (blue) to “GGC” (Gly) in the +1 frame (orange), with the remainder continuing in the zero frame (blue) (Fig. 4B *iii*).

### Luciferase reporters demonstrate efficient +1 frameshifting with *YFS1* frameshift cassette

To investigate if the putative frameshift can support ribosome frameshifting *in vivo* we inserted a 60-nt sequence spanning the putative frameshift site of *YFS1* and flanking regions-7 nt upstream and 46 nt downstream – and inserted in in the SalI/BamHI sites of the pJD375 dual luciferase reporter. In addition, reporter that has the shift site mutated from CUU-AGG-C to CUA-AGG-C and another with the wild-type Ty3 frameshift site were also generated. The plasmids were transformed in BY4741 *S. cerevisiae* strain and the luciferase activity was assayed. When compared to in-frame controls in which Renilla and firefly luciferases are encoded in the same reading frame, these reporters generated 48%, 1.5% and 4.5% frameshifting respectively (Fig. 5).

**Figure 5.**
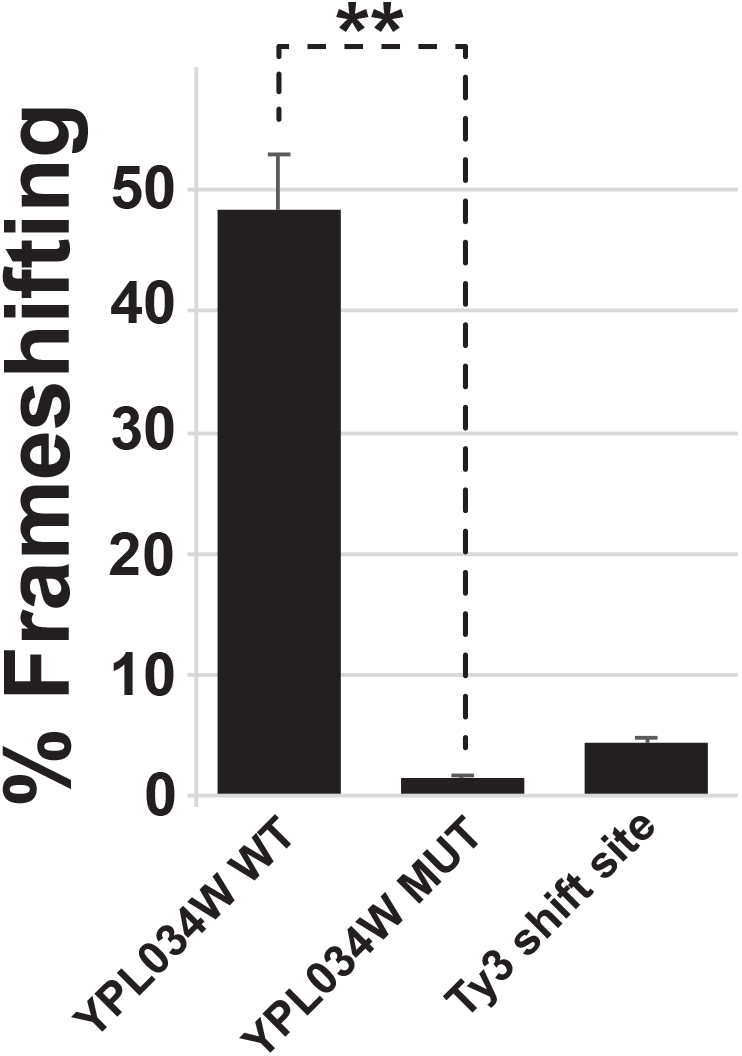
A *YFS1* frameshift cassette supports efficient +1 frameshifting in *S. cerevisiae*. *S. cerevisiae* cells, strain BY4741, were transformed with dual-luciferase reporters containing either wild-type *YFS1* frameshift cassette (60 nt surrounding the putative shift site), a mutant cassette in which the *YFS1* shift site is changed from CUU-AGG-C to CUA-AGG-C, or Ty3 frameshift sequence. In all cases the firefly to *Renilla* ratios are compared to in-frame control values and results are plotted as percent frameshifting. ** p<0.01 (Student’s two-tailed t test; n = 3).

## Discussion

Programmed frameshifting is a well-recognized means of translational regulation of gene expression (Farabaugh 1996; Atkins et al. 2016). While occurring frequently in viruses and retrotransposons (Farabaugh 1996; Atkins et al. 2016), the number of chromosomal genes whose expression is known to require programmed frameshifting is limited (Atkins et al. 2016). Here we describe the identification of only the fourth example of programmed frameshifting in a chromosomally encoded gene in *S. cerevisiae*, which we call *YFS1*, encompassing the previously identified ORF *YPL034W*.

The hypothesis that *YFS1* requires +1 frameshifting for expression is supported by several lines of evidence. The amino acid conservation in the *YPL034W* ORF extends 300 nt upstream of its annotated AUG start codon but this conserved N-terminal extension lacks its own in-frame initiation codon. An upstream ORF is present in all homologs of *YPL034W* from the genus *Saccharomyces*, is always present in the −1 frame relative to *YPL034W*, and overlaps its conserved N-terminal extension. This uORF contains the conserved sequence CUU-AGG-C that is known to be sufficient to direct very efficient +1 frameshifting in *S. cerevisiae* and serves as the +1 frameshift site in the *gag-pol* gene of the retrotransposon Ty1 and the chromosomal *ABP140* gene in the same related organisms (Farabaugh et al. 2006). Aggregate 80S ribosome profiling data (Michel et al. 2014) show continuous uninterrupted ribosome occupancy in the gap between the uORF and the annotated start codon of *YPL034W*. Reading frame analysis indicates that translation of this region is in the +1 frame relative to the uORF and hence the same frame as *YPL034W*. The same data also suggest that the change in reading frame begins at the exact position of the CUU-AGG-C sequence, with the ribosome A site switching from the AGG Arg codon in the 0 frame to the GGC Gly codon in the +1 frame. The first published 40S ribosome profiling data (Archer et al. 2016) do not provide evidence for initiation at the annotated start codon of *YPL034W*, while showing clear evidence of initiation at the AUG of ORF1. Finally, experiments with luciferase reporters (Fig. 5) show that the putative shift sequence of *YFS1* supports robust, 48%, translational +1 frameshifting. This frameshifting level is even higher than the frameshifting inferred from the ribosome profiling data. The difference could be due to different strains used, different growth conditions or sequences outside those tested that suppress the frameshifting in the endogenous locus.

Both the N-terminal extension of *YPL034W* and the requirement for +1 frameshifting for its expression are ancient, emerging perhaps 100-150 million years ago. The N-terminal extension is older and occurs in all homologs of *YPL034W*, even those where it is encoded in the zero frame and can be translated without frameshifting. At least one of the four residues in ORF1 prior to the frameshift, Cys3 in *S. cerevisiae YFS1*, is conserved in orthologs of the gene lacking frameshifting, suggesting that ORF1 is not simply there to provide initiation of translation, but also encodes functional amino acids crucial for the protein’s function.

Farabaugh et al., (Farabaugh et al. 2006) showed that there is a strong correlation between the presence or absence of certain 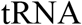 species capable of decoding the CUU Leu codon of the frameshift site and the presence or absence of +1 frameshifting in the *ABP140* and *EST3* genes. Specifically, 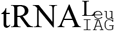 is almost always absent in organisms with +1 frameshifting in *ABP140* and *EST3*, but present in organisms that express these genes without frameshifting, whereas 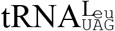 is abundant in organisms with +1 frameshifting in *ABP140* and *EST3* and scarce in organisms without +1 frameshifting in these genes. Ivanov et al. (Ivanov et al. 2006), likewise, found a correlation between the identity of the P-site codon during yeast antizyme +1 frameshifting and the absence of certain tRNA species in specific branches of yeast evolution. This is consistent with the idea that near-cognate tRNA in the P site, in combination with a scarce A-site codon, is a driving force behind efficient +1 frameshifting, as first proposed by Hansen et al (Hansen et al. 2003). The 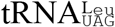 that most frequently decodes the CUU Leu codon in *S. cerevisiae* is unusual in that the uridine in the wobble position of the anticodon is not modified, unlike in more distant yeast relatives where modification of the U restricts it to reading A or both A and G. As a result, 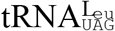 can decode all six Leu codons in *S. cerevisiae* (Randerath et al. 1979; Johansson et al. 2008). Relatively weak pairing of U•U in the wobble position appears to make 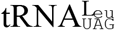 unusually prone to frameshifting when a CUU codon is present in the P site. Work by Vimaladithan and Farabaugh (Vimaladithan and Farabaugh 1994) showed that 11 P-site codons, including CUU, can support appreciable levels of +1 frameshifting. Overexpression of the cognate tRNA for seven of these codons reduced frameshifting, suggesting that frameshifting occurs when the P-site codon is occupied by near-cognate tRNA (Sundararajan et al. 1999). The other four codons lack a predicted cognate decoder. This includes the GCG Ala codon present in the P site of yeast antizyme. Expressing synthetic cognate tRNAs for these four codons reduced frameshifting (Sundararajan et al. 1999), again consistent with the idea that frameshifting occurs when the P-site codon is occupied by near-cognate tRNA. Unlike “simultaneous slippage” −1 frameshifting events, where strong base-pairing between the tRNA and mRNA in the new frame is essential (Weiss et al. 1987; Jacks et al. 1988; Atkins et al. 2016), strong base-pairing is not crucial for P-site driven +1 frameshifting (Hansen et al. 2003; Herr et al. 2004). In other words, it appears that +1 frameshifting is triggered by suboptimal base-pairing of the tRNA in the P site, but once the tRNA is mobilized good base-pairing in the new frame is not critically important.

The emergence of +1 frameshifting in *YFS1* during evolution matches almost perfectly the emergence of the related +1 frameshifting sites in the yeast genes *ABP140* and *EST3*. In those cases as well, no species prior to the evolutionary divergence between <u>*Kluyveromyces*</u> and other *Saccharomycetaceae* shows a requirement for +1 frameshifting for expressing *ABP140* and *EST3* (Farabaugh et al. 2006). This evolutionary pattern matches the chronology of loss of the wobble uridine modifying enzyme for the major CUU-decoding 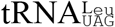., which, as outlined above, appears to drive the shiftiness of CUU-A in *S. cerevisiae*, thus emphasizing the intimate link between the two events.

In addition to the shift-prone codon in the P site, another determining factor for the efficiency of +1 frameshifting is the slow speed of decoding of the zero frame A-site codon. This was first demonstrated for bacterial release factor 2 (RF2). The A site codon of *RF2* during +1 frameshifting is a UGA stop codon, which is recognized uniquely by RF2. Low abundance of RF2 leads to slow decoding of the A-site codon and increased frameshifting, resulting in greater synthesis of RF2, closing an autoregulatory loop (Craigen et al. 1985; Craigen and Caskey 1986; Adamski et al. 1993). Slow decoding of the A site is also important for the efficiency of antizyme +1 frameshifting (Karamysheva et al. 2003). The zero frame A site of Ty1 *gag-pol, ABP140* and *YFS1* is occupied by the AGG Arg codon, which is scarce in *S. cerevisiae*, and the scarcity of the cognate 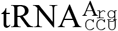 is important for high levels of frameshifting. Overexpression of 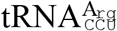 reduces frameshifting, suggesting that slow decoding of the A site is important for efficient frameshifting on the sequence CUU-AGG-C (Belcourt and Farabaugh 1990). Although AGG is highly conserved at the A site of *YFS1* frameshift sites, three species in the *Kazachstania* genus have CGG in place of AGG. CGG also encodes Arg and is used even less frequently than AGG, at 0.8 per 1000 codons versus 3.9 for AGG in *S. cerevisiae* (Nakamura et al. 2000), and is the rarest sense codon for any amino acid in *Kazachstania exigua*. This perhaps explains why it is selected for the A site of some *YFS1* shift sites in this yeast genus.

The role of the last, 7^th^ nucleotide of the heptanucleotide sequence CUU-AGG-C in stimulating +1 frameshifting is only partially understood. Pande et al. (Pande et al. 1995) showed that changing this nucleotide to any one of the other three severely reduces frameshifting. In the same study the authors showed that in an engineered artificial shift site, where the +1 A site is a rare codon, overexpression of the cognate tRNA for the rare codon led to increased +1 frameshifting, seemingly “pulling” the ribosome out of frame. GGC is indeed a common codon in *S. cerevisiae*, with its abundant cognate 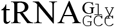 perhaps helping to trap the ribosome in the +1 frame by capture of the new A site following mobilization of the tRNA in the P site. Other explanations for the function of the 7^th^ nucleotide are also possible. Changing the 7^th^ nucleotide of the *S. cerevisiae* OAZ1 +1 frameshift heptanucleotide sequence, GCG-UGA-C, to any other nucleotide severely reduces frameshifting even though all four GAN codons are common, ruling out the abundance of their tRNAs as the determining factor in that case. By contrast, changing the 7^th^ nucleotide of the *S. cerevisiae* Ty3 +1 frameshift heptanucleotide sequence to any of the other three did not reduce frameshifting substantially (Pande et al. 1995). In the case of *S. cerevisiae* OAZ1 the 7^th^ nucleotide is believed to provide unfavorable context for recognition of the UGA stop codon in the A site (McCaughan et al. 1995; Ivanov et al. 2006). It is currently unknown if the “C” in the 7^th^ position of the *YFS1* frameshift site serves a “context” function in the decoding of the adjacent AGG codon in the A site or is there to ensure that an abundant codon is present in the +1 A site.

Occurrence of shift-prone sequences randomly in a coding region would in most cases result in production of truncated polypeptides. Apart from being wasteful, some of these truncated polypeptides could have dominant negative activity that would interfere with the function of the full-length product. This will be especially true for the high frequencies of spurious frameshifting that would be afforded by programmed frameshifting sites. Accordingly, such sequences should be under strong purifying selection in organisms where they occur. Indeed, bioinformatic analysis on coding sequences from *S. cerevisiae* showed that a number of heptanucleotide sequences are under strong negative selection when present in-frame, even when amino acid pairing combinations and codon usage are taken into account. CUU-AGG-C is among the heptanucleotides most highly selected against, as are several of its variants starting with the sequence CUU-AG, five of which score among the 25 most underrepresented heptanucleotides. When tested individually they supported high-level (>5%) +1 frameshifting (Shah et al. 2002). This suggests that there is an undiscovered fundamental feature of this sequence, beyond the unusual tRNA in the P site or the rare Arg codon in the A site that makes it prone to +1 ribosome frameshift.

The *YPL034W/YFS1* gene is non-essential in *S. cerevisiae* and no functions have been assigned to the protein. The closest homolog of Yfs1 that can be identified in BLAST searches is the *S. pombe* protein mug113 (SPAC3F10.05c). The *mug113* gene was identified in a screen for genes whose expression was upregulated during meiosis (Mata et al. 2002; Martin-Castellanos et al. 2005); however, no meiotic phenotypes were observed upon deleting *mug113* in *S. pombe*. Like mug113, the central domain of the YPL034w/Yfs1 protein shows sequence similarity to the GIY-YIG nuclease family. These nucleases are involved in many cellular processes related to DNA repair and recombination including transposon mobility (Mitchell et al. 2019). Consistent with a function related to DNA repair, synthetic genetic array studies (Usaj et al. 2017) revealed that cells lacking *YPL034W* showed similar genetic interactions as cells lacking the Holliday junction resolvase *YEN1* with both genes showing strong synthetic growth defects when deleted in combination with *POL12* encoding the B subunit of the DNA polymerase α-primase complex. As *POL12* is required for mitotic and premeiotic DNA replication, this genetic linkage of *YPL034W* to *POL12* might also relate to the meiotic function of the *mug113* gene in *S. pombe*. As *YPL034W* shows the strongest genetic interactions with loss of the RNA polymerase III subunit *TFC6*, which does not show genetic interactions with *YEN1*, additional studies will be needed to definitively reveal the function of the Yfs1 protein and to determine whether it possesses nuclease activity or functions during meiosis in budding yeast. It will also be of interest to learn whether the utilization of +1 frameshifting for expression of *YFS1* serves a regulatory purpose.

## Materials and Methods

### Sequence Compilation and Analysis

Sequences were obtained from GenBank using BLAST algorithm (Altschul et al. 1990) and applying a strategy described previously (Ivanov et al. 2010) by querying the Nucleotide collection (nr/nt), the RefSeq Genome Database (refseq_genomes) and Whole-genome shotgun contigs (wgs) databases. Sequences were aligned using ClustalX (Larkin et al. 2007). The phylogenetic figure was drawn using the browser based tool iTOL (https://itol.embl.de/) (Letunic and Bork 2019). Logogram figures were generated using WebLogo browser tool (https://weblogo.berkeley.edu/logo.cgi) (Crooks et al. 2004).

### Analysis of Ribosome Profiling Data

The ribosome profiling data were analyzed using the web browser based RiboGalaxy tools (Michel et al. 2016). Briefly, reads from wild type *S. cerevisiae* ribosome profiling data from Guydosh and Green 2014 (Guydosh and Green 2014), Guydosh and Green 2017 (Guydosh and Green 2017), Young et al. 2015 (Young et al. 2015), Young et al. 2018 (Young et al. 2018), and Santos et al. 2019 (Santos et al. 2019) were aligned to the specified mRNA transcript (Ty1 *gag-pol* or *YFS1*) using Bowtie. Up to 2 valid alignments were allowed per read. The minimum seed length was set at 25 nt. CSV files were generated with the reads coming from fragments of 28 nt mapped to each gene with the 5’ end offset from the A site by 15 nt. These files, with the fragment counts per nucleotide position, were used to calculate coverage for each reading frame and to generate figures for that coverage.

### Plasmid Construction

The plasmids listed in Table S3 were constructed by annealing oligos listed in Table S2 bearing the respected frameshift sequences with Sal1/BamH1 overhangs were cloned into pJD375 (ref??) at Sal1/BamH1sites.

### Dual Luciferase Assay

Strain BY4741 (MATa his3Δ1 leu2Δ0 met15Δ0 ura3Δ0) (Research genetics) were transformed with the appropriate constructs or with the corresponding empty vectors (Table S1). To assay luciferase reporters, transformed cells were grown in SC-ura liquid media to an O.D.600nm of 0.8. 1.5ml of exponentially growing cells were lysed with glass beads in 400ul of ice-cold lysis buffer (1× PBS containing two Complete EDTA-free Protease Inhibitor Cocktail Tablets (Roche)/50 mL). Renilla and firefly luciferase activities were measured using 5 μL of 1:10 diluted lysate using the Dual-Luciferase Reporter Assay System (Promega) as described by Dyer et al. (2000) with a small modification in the Stop and Glo substrate (the homemade Stop and Glo buffer was supplemented with 5% commercial Stop and Glo buffer purchased from Promega). CentroXS3 LB960 microplate luminometer fitted with two injectors was used to measure the relative light units (Berthold Technologies). For the YPL034W and Ty3 dual luciferase reporters, Renilla was normalized to firefly luciferase made downstream of the same cistron and the values of “wild type” were compared to “in-frame” control to calculate percent frameshifting. Whereas, in case of the Ty1 dual luciferase reporters, Renilla was normalized to firefly luciferase made downstream of the same cistron and the values of “wild type” were compared to “empty vector” to calculate percent frameshifting.

### Quantification and Statistical Analysis

Luciferase assay data is presented as the mean and standard deviation from biological replicates indicated in the figure legends. The statistical significance between groups and comparisons were made using Student’s two-tailed t tests, using the formula imbedded in Microsoft Excel. p values less than 0.05 were considered significant.

---

**Table S1.** Plasmids used in this study.

| Plasmid | Description | Source/Reference |
| --- | --- | --- |
| YPL034W_WT | YPL034W frameshift site in pJD375 vector | This study |
| YPL034W_MUT | YPL034W mutated frameshift site in pJD375 vector | This study |
| YPL034W_IF | YPL034W inframe frameshift site in pJD375 vector | This study |
| Ty3_WT | Ty3 frameshift site in pJD375 vector | This study |
| Ty3_IF | Ty3 inframe frameshift site in pJD375 vector | This study |
| Ty1 | Ty1 frameshift site in pJD375 vector | ref |
| pJD375 | empty vector | ref |

**Table S2.**
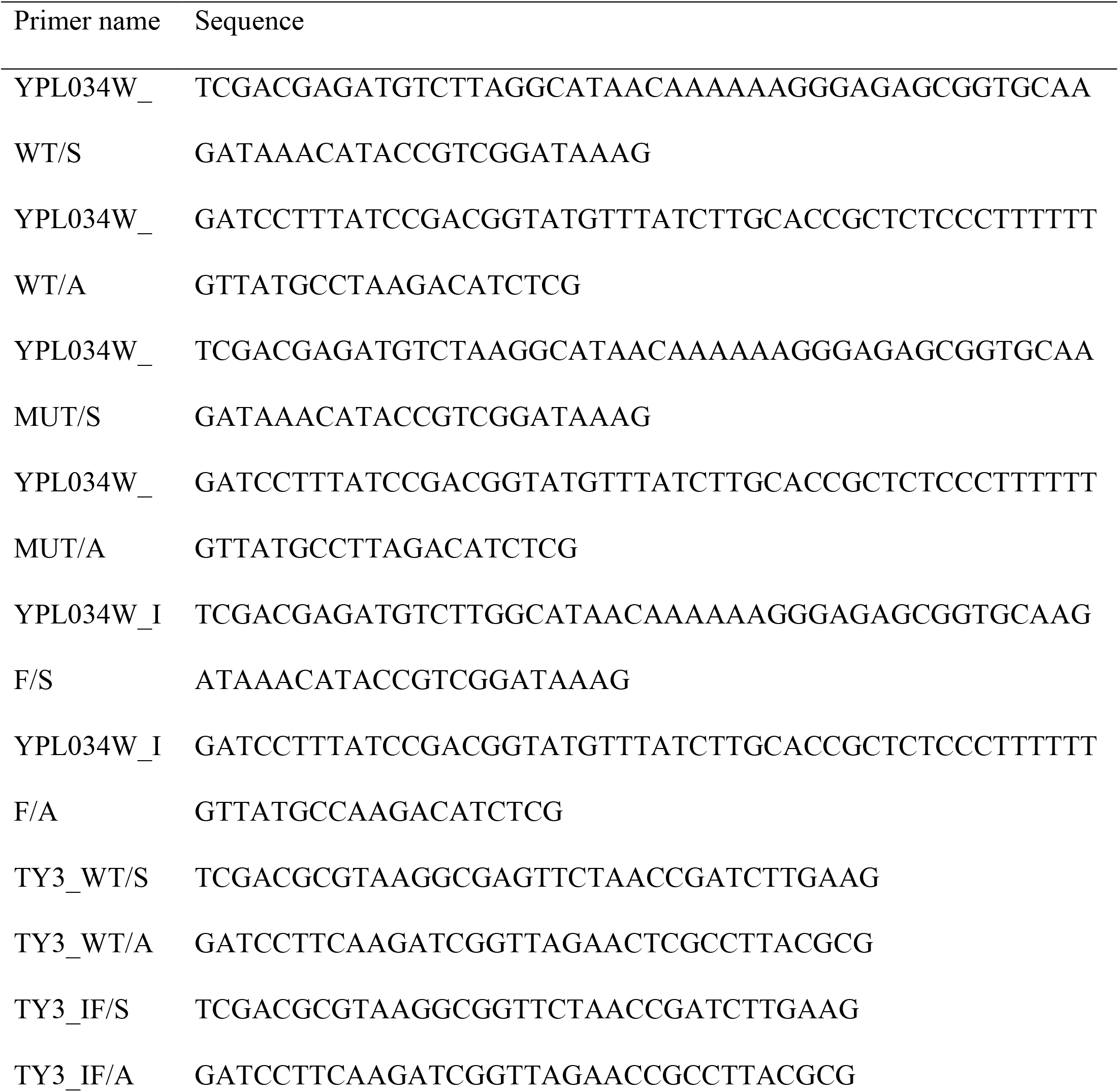
Oligonucleotides used in this study.

**Table S3.**
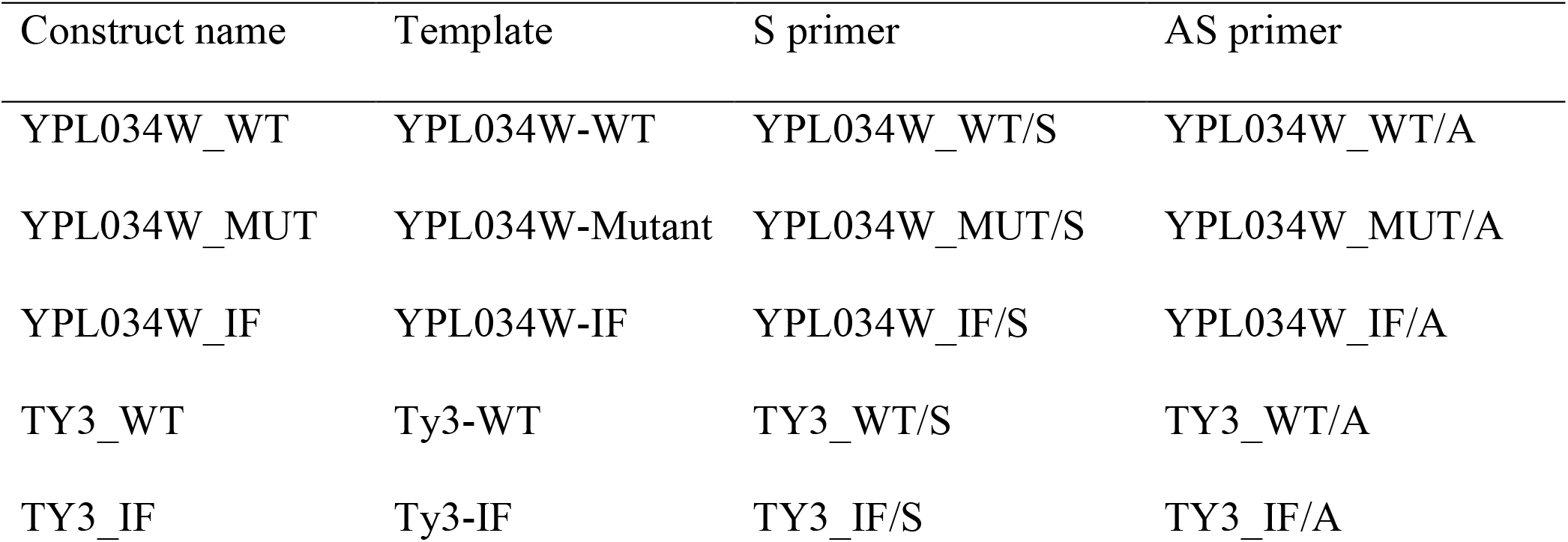
Plasmids made for this study.

| Construct name | Template | S primer | AS primer |
| --- | --- | --- | --- |
| YPL034W_WT | YPL034W-WT | YPL034W_WT/S | YPL034W_WT/A |
| YPL034W_MUT | YPL034W-Mutant | YPL034W_MUT/S | YPL034W_MUT/A |
| YPL034W_IF | YPL034W-IF | YPL034W_IF/S | YPL034W_IF/A |
| TY3_WT | Ty3-WT | TY3_WT/S | TY3_WT/A |
| TY3_IF | Ty3-IF | TY3_IF/S | TY3_IF/A |

